# Arabidopsis PRMT5 Buffers Pre-mRNA Splicing and Development Against Genetic Variation in Donor Splice Sites

**DOI:** 10.1101/2024.11.26.624665

**Authors:** Maximiliano S. Beckel, Abril San Martín, Sabrina E. Sánchez, Danelle K. Seymour, María José de Leone, Daniel Careno, Santiago Mora-García, Detlef Weigel, Marcelo J. Yanovsky, Ariel Chernomoretz

## Abstract

Genetic variation at splice site signals significantly influences alternative splicing, leading to transcriptomic and proteomic diversity that enhances phenotypic plasticity and adaptation. However, novel splice variants can negatively impact gene expression and developmental stability. Canalization—the ability of an organism to maintain a consistent phenotype despite genetic or environmental variations—helps balance the effects of genetic variation on development and evolution. Protein arginine methyltransferase 5 (PRMT5) is a key splicing regulator in plants and animals. Most splicing changes in *prmt5* mutants are linked to weak donor splice sites, suggesting that PRMT5 may buffer splicing against genetic variation. We examined PRMT5’s effects on splicing and development in two genetically divergent *Arabidopsis thaliana* accessions with different single nucleotide polymorphisms (SNPs) affecting donor splice sites. While PRMT5 inactivation similarly affected splicing in both backgrounds, it significantly increased splicing and phenotypic differences between the accessions. Our findings suggest that PRMT5 contributes to canalization, mitigating the impact of splice site polymorphisms and facilitating the evolution of adaptive splicing patterns.

## INTRODUCTION

Deciphering how genetic variation influences gene expression and phenotypic diversity is essential for understanding evolutionary processes and ecological adaptations (Wright, Smith, and Jiggins 2022). Variations in gene expression can stem from genetic differences that impact transcription factor protein dynamics or from changes in DNA sequences associated with regulatory elements such as enhancers or promoters. Alterations in co-or post-transcriptional regulatory mechanisms also contribute to these variations (Wright, Smith, and Jiggins 2022). In eukaryotes, most genes include non-coding DNA sequences called introns, which are removed from pre-mRNA during the splicing process (Kornblihtt et al. 2013). The accurate removal of introns and the joining of exons—segments of a gene retained in the mature messenger RNA—are critical for proper gene expression. During splicing, different exons can be combined in various ways, such as by skipping one or more exons, retaining an intron, or utilizing alternative donor or acceptor splice sites. This phenomenon, known as alternative splicing (AS), enables the generation of diverse transcripts from a single gene, thereby enhancing variability in both the transcriptome and proteome, which may also contribute to regulating transcript levels (Kornblihtt et al. 2013). AS and its regulation are crucial for numerous biological processes, including physiological and developmental functions in both plants and animals (Marasco and Kornblihtt 2023; Staiger and Brown 2013). Comparative studies have demonstrated that AS evolves more rapidly and is more species-specific than other transcript-level changes, contributing to phenotypic differences among organisms and highlighting AS as a key mechanism driving evolutionary diversity (Verta and Jacobs 2022).

Pre-mRNA splicing can be influenced by mutations in core or auxiliary splicing factors or, more commonly, by mutations in sequence motifs found near intron-exon junctions such as core 5’ splice sites (5’ss), 3’ splice sites (3’ss), branch point sequences, or splicing regulatory elements (SREs) (G.-S. Wang and Cooper 2007). While genetic variation that alters pre-mRNA splicing can lead to evolutionary innovations, it can also disrupt gene expression or function, negatively impacting organismal performance. Indeed, genetic variation affecting pre-mRNA splicing has been associated with human genetic diseases (Rogalska, Vivori, and Valcárcel 2023). Specifically, mutations in 5’ss frequently lead to intron retention, exon skipping, or other significant alterations to the mRNA and the encoded protein (De Conti et al. 2012). Consequently, buffering or compensatory mechanisms probably exist to mitigate the potentially detrimental effects of such mutations while allowing genetic variation at 5’ss to accumulate and contribute to evolutionary innovations (Rutherford 2000). This hypothetical buffering mechanism would be analogous to the role of Heat Shock Protein 90 (HSP90) as a capacitor that buffers the effects of genetic variation on protein function by preventing the aggregation of misfolded proteins (Rutherford and Lindquist 1998; Queitsch, Sangster, and Lindquist 2002). This buffering capacity allows organisms to exhibit stable traits despite underlying genetic diversity.

Although progress has been made in understanding how genetic sequences acting in *cis* (i.e., located near the focal gene) buffer the effects of weak 5’ss on pre-mRNA splicing (Xiao et al. 2009), the role of *trans*-acting factors (e.g., diffusible regulatory proteins encoded elsewhere in the genome) in alleviating the negative consequences of mutations affecting 5’ss strength remains largely unknown.

Protein Arginine Methyltransferase 5 (PRMT5), a type II PRMT responsible for symmetric dimethylation of arginine residues in various histone and non-histone proteins, has been identified as a major regulator of pre-mRNA splicing in plants and animals (Sanchez et al. 2010; Deng et al. 2010; Bezzi et al. 2013). Among other proteins, PRMT5 methylates three of the seven Sm proteins (D1, D3, and B), which are core components of U1-U5 snRNPs (Meister et al. 2001); this methylation is vital for the correct maturation of these spliceosomal complexes in human cells (Gonsalvez et al. 2007). Furthermore, PRMT5 has also been shown to methylate the LSm4 protein, a core component of U6 snRNP, in both plants (Agrofoglio et al. 2024) and humans (Arribas-Layton et al. 2016).

Functional impairment of PRMT5 has been linked to various phenotypes. In plants, *prmt5* mutants show alterations in the floral transition and other developmental processes, circadian rhythms, and responses to salt stress (Sanchez et al. 2010; Hong et al. 2010; Z. Zhang et al. 2011; Pei et al. 2007). In mammals, PRMT5 influences the proper splicing of the gene encoding the MDM4 protein, a crucial suppressor of the p53 pathway (Bezzi et al. 2013). Thus, PRMT5 holds significant clinical relevance for researching and treating various types of cancer (Rengasamy et al. 2017; Radzisheuskaya et al. 2019; Kim and Ronai 2020; Sachamitr et al. 2021).

Studies in plants and animals indicate that PRMT5 modulates splicing of constitutive as well as AS events associated with weak donor splice sites, which deviate from the consensus sequence (Sanchez et al. 2010; Bezzi et al. 2013; Hernando et al. 2015). While its effects on AS led us to propose that PRMT5 helps organisms to synchronize developmental and physiological processes with changes in environmental conditions ((Sanchez et al. 2010; Hong et al. 2010; Z. Zhang et al. 2011)), the functional relevance of its impact on constitutive splicing is less clear. Here we propose that PRMT5 also functions as a key component of a buffering mechanism that attenuates the effect of natural genetic variation at 5’ss, allowing organisms to maintain a high frequency of 5’ss polymorphism that may facilitate the evolution of new splicing patterns. To evaluate this hypothesis we compared the impact on pre-mRNA splicing of mutating the *PRMT5* gene in two different genetic backgrounds (i.e. different accessions) of *Arabidopsis thaliana*. We found strong evidence supporting the idea that PRMT5 plays a key role in maintaining splicing efficiency amid genomic variations in donor splice sites, potentially acting as a safeguard for system stability. This work highlights the value of testing mutations in different genetic backgrounds to better understand the complex interplay between gene variants, genomic background, and phenotypic diversity (Srikant et al. 2022; Koneru et al. 2021), (Milloz et al. 2008).

## MATERIALS AND METHODS

### Plant material and growth conditions

The Columbia (Col-0) and Landsberg *erecta* (Ler) accessions of *Arabidopsis thaliana* (*A. thaliana*) were employed as wild types (WT) for physiological assays and as the genetic background for *prmt5* mutations. Plants were cultivated on soil at 22°C under long-day (LD; 16-h light/8-h dark cycles; 80 μmol·m^-2^ s^-1^ of white light) conditions.

### Introgression of the *prmt5-5* into a different genomic background

The *prmt5-5* mutant isolated in the Col-0 background was backcrossed for 7 generations with WT plants of the L*er* accession, checking for the presence of the mutant allele after each cross. To generate hybrid backgrounds of WT plants (F1 Col-0 x L*er*) and *prmt5-5* mutant plants (F1 *prmt5-5* Col x *prmt5-5* L*er*), multiple crosses were conducted involving WT and *prmt5-5* mutant plants from the Col-0 and L*er* accessions. The resulting seeds and seedlings were analyzed for transcriptome characterization using RNA-seq.

### Novel *prmt5* mutant alleles in Col-0 and L*er* accessions by CRISPR-Cas9 technology

Independent *prmt5* mutant lines (*prmt5-6*) were generated in the Col-0 and L*er* accessions by using CRISPR/Cas9 technology. Two single guide RNAs (sgRNAs) were designed for the *PRMT5* gene using the CCTOP web tool (Stemmer et al. 2015) and incorporated in forward and reverse PCR primers, respectively, as previously described by Xieng et al (Xing et al. 2014) (see Supplementary Table S1). The PCR fragment was amplified from the pCBC-DT1T2 plasmid using Q5 DNA polymerase (New England Biolabs, NEB). The insert was then cloned into the binary vector pHEE401E via the Golden Gate cloning method, as previously described by Xieng et al (Xing et al. 2014). The resulting plasmid was then transformed into Col-0 and L*er* plants using the *Agrobacterium tumefaciens* GV301 strain with the floral dip method (X. Zhang et al. 2006). Transformed seedlings were selected on Petri plates containing Murashige and Skoog (MS) medium 0.8% m/v of agar and hygromycin (20 mg/L). Homozygous mutants were identified by amplifying the sequences around the sgRNA target sites with specific primers followed by sequencing by capillary electrophoresis sequencing (CES), which revealed a deletion of 301 bp relative to the WT (see Fig. S1). Transgene-free homozygous mutants were selected by PCR diagnosis.

### Flowering Time Analysis

For flowering time measurements, plants were grown on soil at 22°C under standard long-day conditions. Flowering time was estimated by counting the number of rosette leaves at bolting. These experiments were performed in quadruplicate with at least 12 individuals for each genotype. The statistical analysis was done using a two-way ANOVA followed by Sidak’s multiple comparisons test with a p-value of 0.05.

### Circadian Leaf Movement Analysis

For leaf movement analysis, WT and *prmt5* mutant plants were entrained under LD conditions until the appearance of the first pair of leaves and then transferred to continuous white light (20 -30 μmol m^-2^ s^-1^) at 22 °C. The position of the first pair of leaves was recorded every 2 h for 5–7 days using digital cameras and the leaf angle was determined using the ImageJ software (http://imagej.nih.gov). The circadian period was estimated using the BioDare2 software (biodare2.ed.ac.uk) (Zieliński, Hay, and Millar 2022) and analyzed with Fast Fourier Transform Non-linear Least Squares (FFT-NLLS) with the same software. The experiments were performed in triplicate with 12 seedlings for each genotype. A two-way ANOVA followed by Sidak’s multiple comparisons test was used to test for statistical significance with an alpha of 0.05.

### RNA extraction and sequencing

Seeds were sown on MS medium containing 0.8% m/v agar, stratified for four days in the dark at 4°C, and then transferred to continuous white light at 22°C. After nine days of growth, entire seedlings were collected and total RNA was purified using the QIAGEN RNeasy Plant Mini Kit following the manufacturer’s protocol. cDNA libraries were generated based on the Illumina TruSeq RNA Sample Preparation Guide. Polyadenylated mRNA was extracted from 3 μg of total RNA and then fragmented. Reverse transcriptase SuperScript II (SSII RT) (Thermo Fisher Scientific, Waltham, MA, USA) and random hexamers were used to synthesize cDNA. Finally, specific adapters were added to each sample and the libraries were sequenced on the Illumina GAIIx platform with 100 bp reads. Reads were analyzed using version 1.3 of the Illumina pipeline and underwent quality filtration using standard Illumina procedures. The resulting sequence files were created in fastq format.

### Mapping of RNA-Seq reads

Before carrying out read mapping, the TrimGalore wrapper was used to remove adapter sequences with Curadapt (Martin 2011) and to check read quality with FastQC. The STAR v2.7.9a (Dobin et al. 2013) aligner was used for read mapping due to its superior performance in aligning reads spanning splice junctions, with only single-mapping reads considered. STAR’s “2-pass” method improved sensitivity in detecting novel junctions that were not aligned to the coordinates of isoforms present in the genome annotation. The TAIR10 reference genome was used for reads from the Col-0 accession, while an established strategy (Winkelmüller et al. 2021) was employed for L*er*. A pseudo-reference genome was generated by utilizing the single nucleotide variant (SNP) and insertion/deletion (Indel) data of L*er* obtained from the 1001 Genomes Project site (https://1001genomes.org/), starting from the Col-0 genome. The pseudogene function incorporated in the GEAN software (Song et al. 2019) was employed to infer the sequence of the pseudo-genome by replacing the reference allele with the alternative allele of L*er*. For the annotation of genetic features, GFF files were constructed by projecting the genomic coordinates from TAIR10 onto the coordinates of the L*er* accession. To achieve this, the liftgff function of GEAN was utilized. Reads from F1 hybrids were aligned using both TAIR10 and the L*er* pseudo-genome. A specific parent was assigned to each read based on which alignment had the highest quality. Only reads that could be unambiguously assigned to one parent were used to analyze alternative allelic splicing. To process the BAM files created from the aforementioned alignments, samtools software tools were utilized (Li et al. 2009). Only properly paired reads with a mapping quality greater than 20 were included in the analysis.

### Differential gene expression and alternative splicing analysis

R package ASpli (Mancini et al. 2021) was used to count the number of overlapping reads with various genomic features, such as genes’ exons/introns or junctions, for the detection of differentially expressed genes (DEG). Likewise, for the differential use of subgenomic regions known as bins in comparisons between wild-type and *prmt5-5* mutants, this package was employed (Mancini et al. 2021). The significance criteria in these comparisons were a fold change (FC) greater than 1.5 and a false discovery rate (FDR) lower than 0.05. To assess for changes in splicing patterns between different accessions a Fisher exact test was considered to quantify the statistical significance between supporting and non-supporting junctions. The resulting p-values were adjusted for multiple comparisons using the BenjaminiHochberg (BH) method. A significance criterion of a q-value less than 0.1 was applied to the three replicates with an average |ΔPIR/PSI| greater than 0.1.

The interaction effect between accession (Col-0 or Ler) and genotype (WT or prmt5 mutant) was estimated using the methodology described by Altman and Bland (Altman and Bland 2003; McManus et al. 2014; Gao et al. 2015; X. Wang et al. 2019). Briefly, log-transformed PSI or PIR ratios between WT and prmt5 mutant plants were calculated for each event in both accessions, with standard errors estimated via bootstrap resampling (Jewell 2003). These were used to compute z-values and corresponding p-values, which were adjusted using the FDR method. A q-value threshold of <0.1 was used to identify significant events.

### Pathways and SNP enrichment analysis

To explore the biological relevance of DEG sets in the WT/*prmt5-5* contrasts, we carried out a KEGG pathway enrichment analysis with the R clusterProfiler package. The BH method was employed in adjusting the p-values with those lower than 0.05 being deemed significant. The R pathways package was utilized to illustrate the metabolic pathways along with the genes differentially expressed within them. The genomic sequences for *A. thaliana* were obtained and manipulated through the use of several R packages, including GenomicFeatures, Biostrings, and rtracklayer. Additionally, the VariantAnnotation R package was used to analyze and manipulate information regarding genomic variations between Col-0 and L*er*.

### Strength assessment of 5’ss sequences

To computationally assess the strength of 5’ss sequences we employed the sequence-based energy estimation method proposed by Beckel and colleagues (Beckel et al. 2023). This quantity, estimated within a maximum-entropy modeling framework, offers a quantitative evaluation of the frequency at which a particular sequence occurs across the entire genome. Sequences with low energy values typically represent highly prevalent 5’ss, whereas those with high energy values tend to correspond to infrequent donor splice sites. This data-driven characterization aligns well with energy scales derived from biochemical dimerization estimations of 5’ss sequences against the U1 RNA stretch (Beckel et al. 2023).

### RNA Isolation and AS Event Validation by RT-PCR Analysis

For alternative splicing validation, three biological replicates of WT and *prmt5* mutants from Col-0 and L*er* accessions were sown on MS-agar plates, cold stratified in darkness for 3 days, and grown for 10 days under LD conditions. Total RNA was extracted using BioZol reagent (Productos Biologicos, PB-L) following the manufacturer’s protocols. To estimate the concentration and quality of the samples, a NanoDrop 2000c (Thermo Scientific, Waltham, MA, USA) and agarose gel electrophoresis were used. One microgram of RNA was treated with RQ1 RNase-Free DNase (Promega, Madison, WI, USA) and subjected to retro-transcription with SuperScript II Reverse Transcriptase (SSII RT) (Thermo Fisher Scientific, Waltham, MA, USA) and oligo-dT according to the manufacturer’s instructions. Amplification of cDNA was carried out using 1.5 U of Taq polymerase (Invitrogen, Carlsbad, CA, USA) for 30-34 cycles to measure the relative abundance of the isoforms at the linear phase of amplification. The primers used for amplification are detailed in Supplementary Table S1. RT–PCR products were electrophoresed in 1% (w/v) agarose and detected by Ethidium Bromide.

## RESULTS

### Experimental design and genetic variation at donor splice site sequences

Trans-acting regulators can influence the effects of genetic background on gene expression, thereby affecting how genetic variation contributes to phenotypic diversity. To investigate how PRMT5, a key transcriptional and post-transcriptional regulator, affects pre-mRNA splicing differences between two *A. thaliana* accessions, we introgressed the *prmt5-5* mutant allele, originally isolated from the Col-0 accession, into the L*er* accession through repeated backcrossing. We then conducted an RNA-Seq experiment to compare the transcriptomes of young WT and *prmt5-5* mutant plants from both Col-0 and L*er* accessions (Fig. 1A).

**Figure 1.**
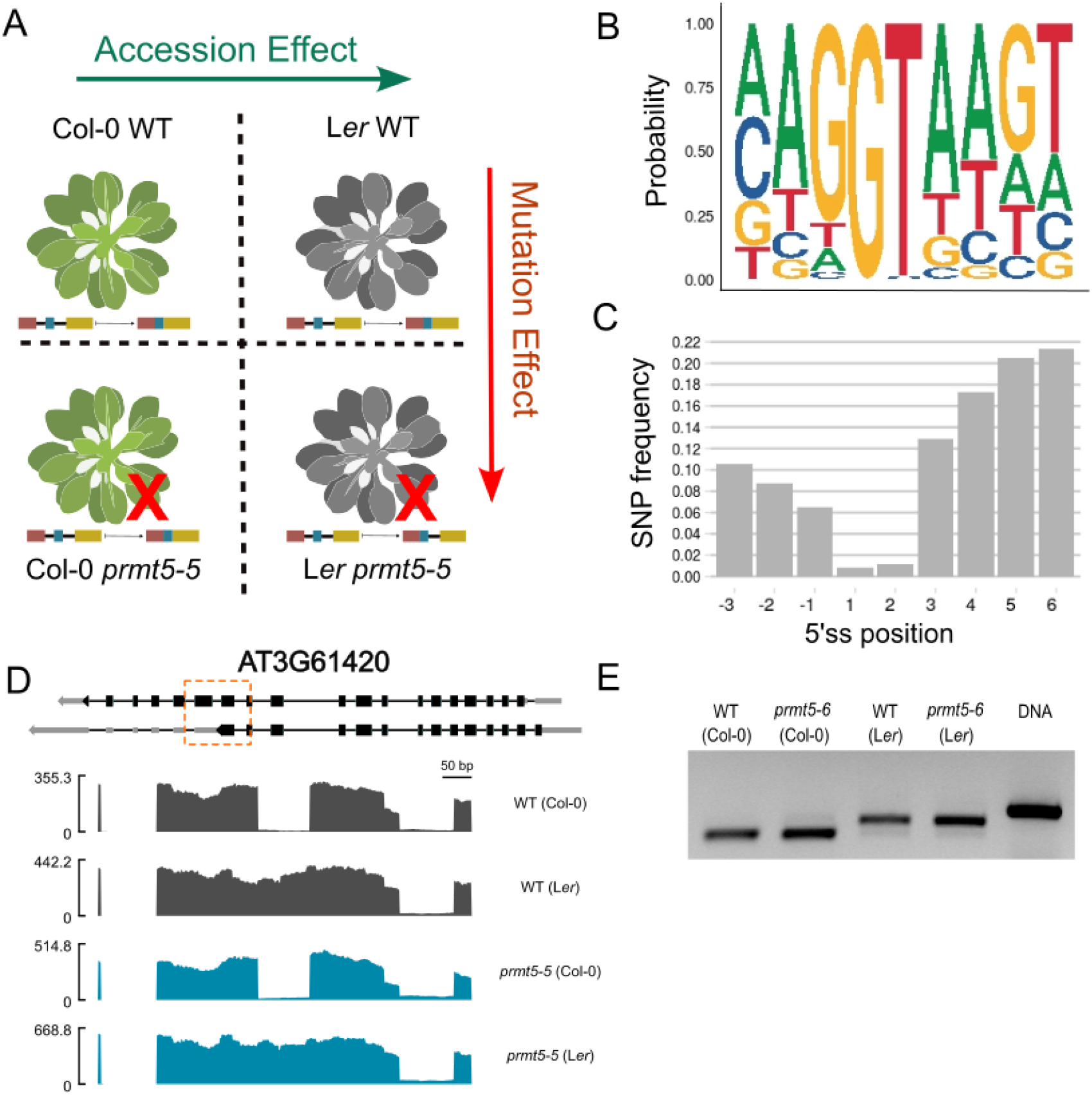
**(A)** Experimental design including WT and *prmt5-5* mutant (red cross in the scheme) plants from two different *A. thaliana* accessions, Col-0 (green plant in the scheme), and L*er* (gray plant in the scheme) (**B**) Logo of 5’ss sequences from Col-0 genome belonging to expressed genes in our samples (153,213 sites). (**C**) Frequency of SNPs between Col-0 and L*er* accessions located in 5’ss sequences (3,615 SNP’s) **(D)** Coverage plot of the AT3G61420 gene in WT (gray) and *prmt5-5* (blue) mutant plants of the Col-0 and L*er* accessions. A representative scheme describing the different isoforms annotated for the gene is shown above the coverage plot. Boxes represent exons, lines represent introns and the orange box surrounds the measured event. **(E)** RT-PCR validation of the differential splicing event shown in panel D, including a genomic DNA control.

Before characterizing the effect of PRMT5 on pre-mRNA splicing in these genomic backgrounds, we analyzed the consensus sequence associated with the donor splice site in *A. thaliana* (Fig. 1B) and determined if there was genetic variation at each position of the splice site between accessions. We found a significant depletion of sequence variation at the +1 and +2 donor splice sites, which is consistent with their critical role in the splicing reaction (Fig. 1C). Reduced genetic variation at these positions indicates that these mutations could be negatively selected, as variation may eliminate splicing at these positions. Indeed, in the few expressed genes with genetic variation at the +1 position of a donor splice site, replacing the consensus nucleotide G at +1 resulted in the complete retention of the associated intron in the accession carrying the non-consensus nucleotide. This pattern was consistent across allelic variants of *PRMT5* present (Fig. 1D,E).

### Common effects of PRMT5 on gene expression and pre-mRNA splicing

We analyzed the impact of the *prmt5-5* mutation on mRNA levels in the Col-0 and L*er* accessions. In our comparison between *prmt5-5* and WT plants, we identified 1,903 differentially expressed genes (DEGs) in the Col-0 accession and 2,508 DEGs in the L*er* accession (Supplementary DataSet). Of these, 975 genes were common to both sets, representing 51.2% of DEGs in Col-0 and 38.9% in L*er*. In both accessions, we observed a greater number of genes with increased expression in the *prmt5-5* mutant compared to the WT. Only a small number of genes were overexpressed in one mutant background but underexpressed in the other background (Fig. S2 A). This confirms that *PRMT5* has similar effects on gene expression in both accessions, acting mostly as a repressor. The consistency in differential gene expression signals between the accessions was also reflected at the level of affected biological processes. A KEGG pathway enrichment analysis revealed that both accessions experienced alterations in a similar set of pathways, primarily related to pre-mRNA splicing regulation and RNA degradation (Fig. S2 B,C). There was also an increase in the expression of genes associated with the main components of the spliceosome in both accessions when comparing the transcriptomes of *prmt5-5* mutants and WT plants. The strong correlation in fold change (R = 0.83) for 80 spliceosome-related genes in Col-0 and L*er* further suggests that PRMT5 has a similar effect in both accessions.

The results of the RNA-seq experiments were then used to characterize changes in pre-mRNA splicing patterns in *prmt5-5* mutants compared to WT samples across both genomic backgrounds. We identified 1.130 differential splicing events between WT and *prmt5-5* mutant plants for Col-0 and 947 for L*er* (SupplementaryDataSet). There was a high concordance across accessions, with 696 shared differential splicing events (61.6% of the differential splicing events in Col-0 and 73.5% in L*er*) (Fig. 2A). Intron retention (IR) was the predominant type of alternative splicing (AS) accounting for 92.1% of events in Col-0 and 94.7% in L*er*.

**Figure 2.**
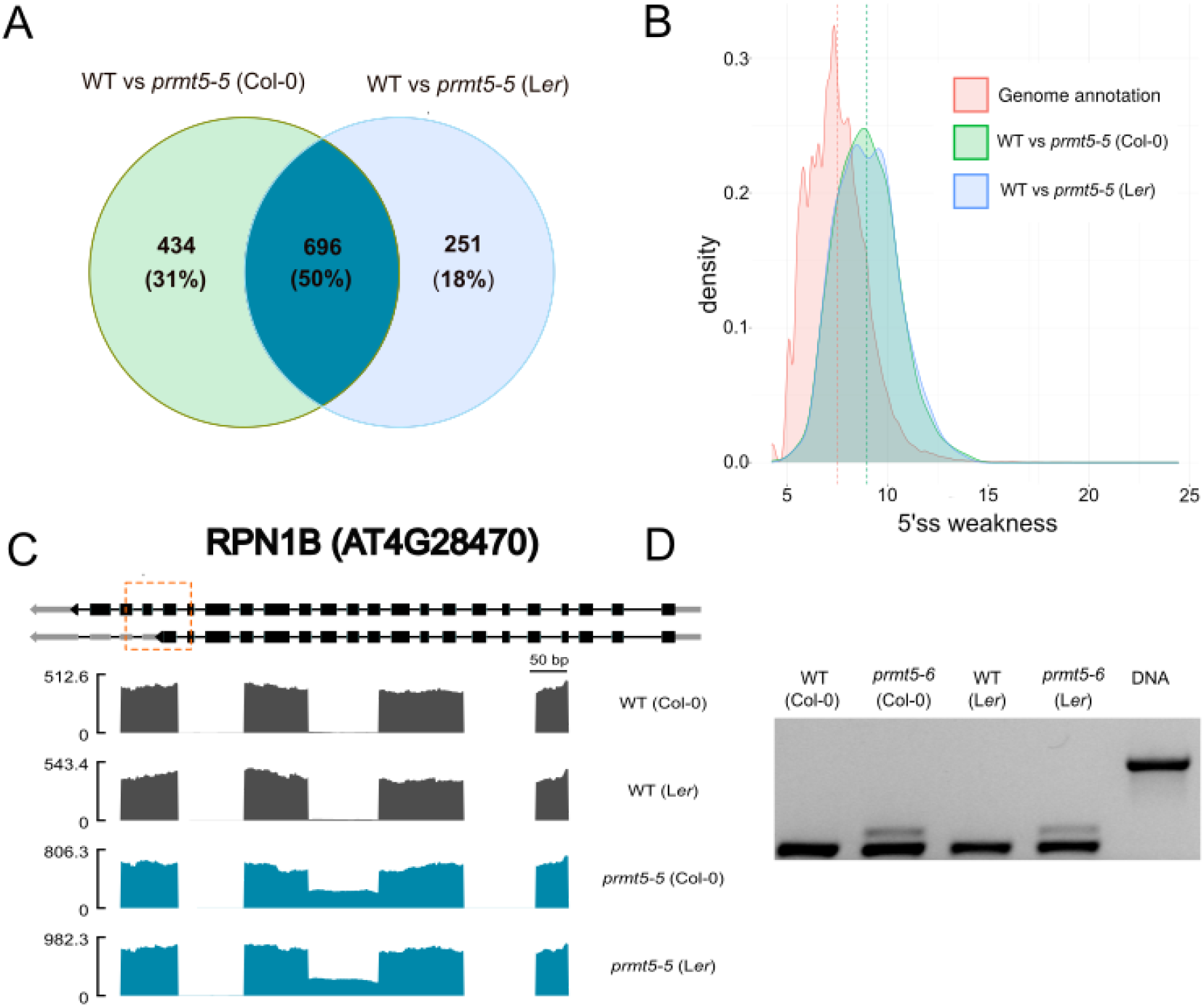
Differential splicing between WT and *prmt5* mutant plants. **(A)** Venn diagram of the bins that underwent a significant change between WT and *prmt5-5* mutant samples in Col-0 and L*er*. **(B)** Distribution of energies for splice donor sites (5’ss) of intron retention events in each of these comparisons. Genome annotation: 5’s extracted from genome annotation and belonging to genes expressed in at least one condition; Col-0 and L*er*: 5’ss from intron retention events in WT vs *prmt5-5* comparison in Col-0 and L*er*, respectively. Dotted lines indicate the mean of the distributions. 5′ss weakness was estimated using a sequence-based energy estimation method (see Materials and Methods). Sequences with low energy values represent highly prevalent (i.e. strong) 5’ss, whereas those with high values correspond to infrequent (i.e. weak) donor splice sites. **(C)** Coverage plot of RPN1B (AT4G28470) in WT (gray) and *prmt5-5* (blue) Col-0 and *Ler* accessions. A representative scheme describing the different isoforms annotated for the gene is shown above the coverage plot. Boxes represent exons, lines represent introns and the orange box surrounds the measured event. **(D)** Validation of the differential splicing event by agarose gel with the RT-PCR amplicons and the corresponding genomic DNA control.

For both accessions, the 5’ss of the differentially regulated splicing events were enriched for non-consensus sequences, supporting a role for PRMT5 in the regulation of splicing events at weaker donor splice sites (Fig. 2B). For the majority of AS events (1,028 out of 1,033 events for Col-0 and 887 out of 893 events for L*er*), there was higher intron retention (as measured by Percent Intron Retention (ΔPIR)) in *prmt5-5* mutants compared WT plants. These results suggest that *prmt5-5* mutant plants in both accessions struggle to recognize weak donor splice sites, leading to increased intron retention. Interestingly, only about 25% of the genes with altered splicing patterns in the *prmt5-5* mutants also showed significant differences in expression levels. This indicates that the effects of PRMT5 on splicing do not depend on overall levels of mRNA. To validate the effect of the *prmt5* mutant allele on pre-mRNA splicing across genomic backgrounds, we conducted semi-quantitative RT-PCR using a novel loss of function deletion allele, *prmt5-6*. This allele, which was generated by editing the *PRMT5* gene in both Col-0 and L*er* accessions using CRISPR/Cas9, displayed pre-mRNA splicing defects similar to those observed in *prmt5-5* mutant plants (Fig. 2C,D).

### PRMT5 effects on differences in splicing between Col-0 and L*er* accessions

Analysis of the differences in splicing patterns between Col-0 and L*er* accessions in WT and *prmt5-5* mutant backgrounds was performed using Fisher’s exact test. We observed 686 and 1,265 differential splicing events between Col-0 and L*er* accessions in WT and *prmt5-5* mutant backgrounds, respectively (SupplementaryDataSet). The difference was mainly due to an increase in the number of intron retention events that differed between Col-0 and L*er* in the *prmt5-5* mutant background (Fig. 3A), indicating that more pronounced differences in splicing patterns were observed between accessions in the absence of *PRMT5*.

**Figure 3.**
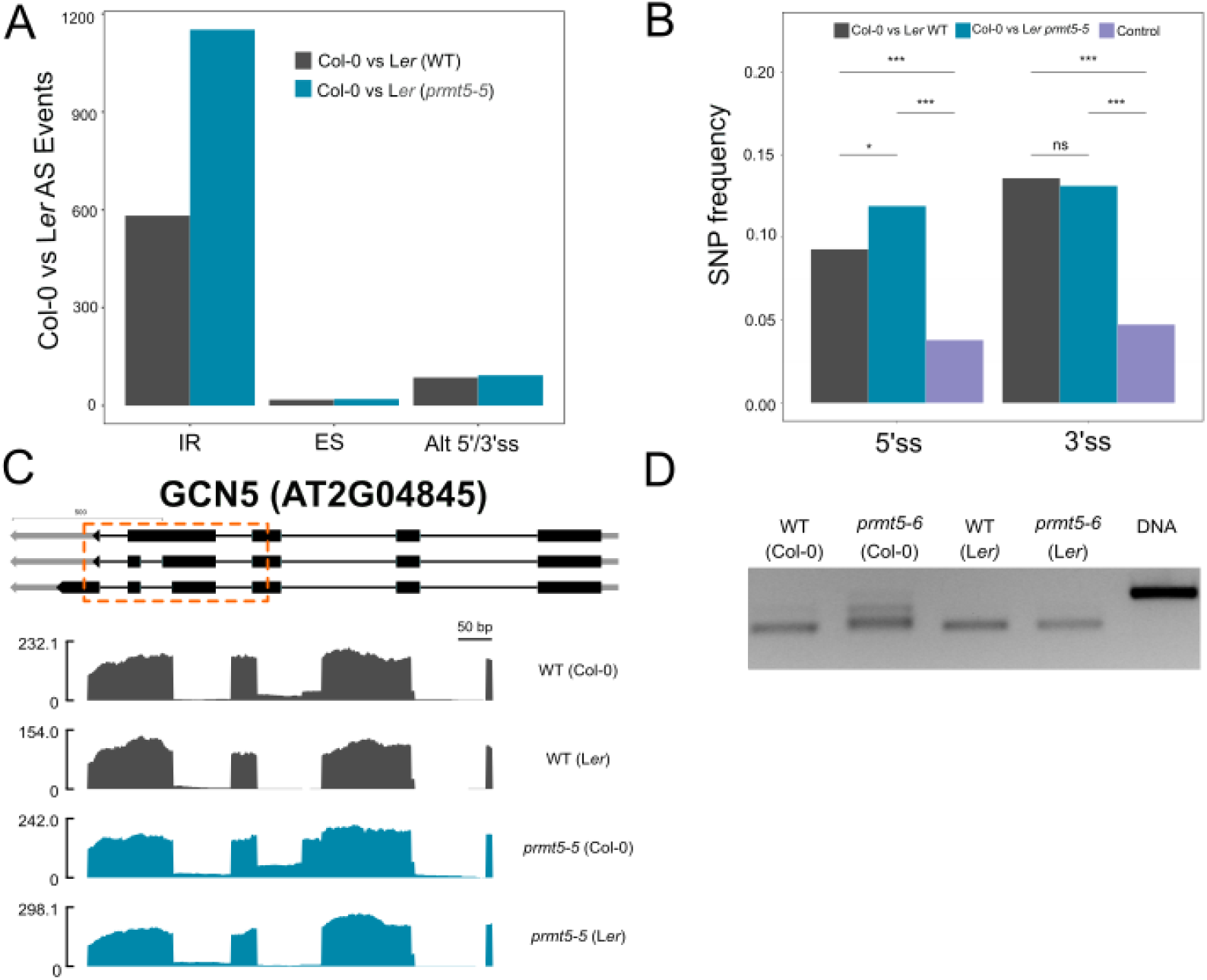
Differential splicing events between Col-0 and L*er*. **(A)** Number of differential splicing events in the comparison between accessions in WT (gray bars) and *prmt5-5* mutant (blue bars) backgrounds according to the type of AS event (IR: intron retention, ES: exon skipping, Alt 5’ss/3’ss). **(B)** Frequency of occurrence of SNPs/indels between Col-0 and L*er* accession at donor (5′ss) and acceptor (3′ss) splice sites for IR events differentially spliced between accessions in WT (gray bars) or *prmt5-5* (blue bars) mutant backgrounds compared to annotated IR events not differentially spliced (Control, violet bars). Fisher’s test to test for differences in observed frequencies. Significance: “***”: p-value < 0.001; “**”: p-value < 0.01; “*”: p-value < 0.05, “ns”: p-value >0.05. **(C)** RNA-seq data coverage plot of the GCN5 (AT2G04845) gene in WT (gray) and *prmt5-5* (blue) mutant plants of the Col-0 and *Ler* accessions. A representative scheme describing the different isoforms annotated for the gene is shown above the coverage plot. Boxes represent exons, lines represent introns and the orange box surrounds the AS event under evaluation. **(d)** Validation of the differential splicing event at the GNC5 gene by RT-PCR.

We sought to assess the possible influence of genetic variation in nearby sequences in determining the intron retention differences observed between accessions in both WT and *prmt5-5* mutant backgrounds. To do this, we calculated the density of SNPs/indels within a genomic region that included both, the retained intron and the two adjacent exons. Statistically significant differences were observed in the density of SNPs/indels in sequences neighboring IR that varied between Col-0 and L*er* and IR events that were shared (see Fig. S3). There was a mean of 6.3 SNPs or indels/1000 bp and 6.7 SNPs or indels /1000 bp surrounding differentially retained introns between accessions in WT and *prmt5-5* mutant plant backgrounds, respectively, which was reduced to 3.5 SNPs or indels/1000bp for annotated IR events that were not differentially spliced between accessions, supporting the idea that the differences in splicing between accessions results from genetic variations in sequences surrounding splice sites (Fig. S3).

We then focused our analysis on sequence variants at 5′ss and 3′ss. For donor splice sites, we considered SNPs located within the last three positions of the exon preceding the retained intron and the first six positions of the intron. For acceptor sites, the last 13 intronic positions and the first position of the following exon were considered. In both cases, we found that the frequency of SNPs was significantly higher for regions that were differentially spliced between accessions, either in WT or *prmt5* mutant backgrounds, compared to regions associated with IR events not differentially spliced between accessions (Fig. 3B). Interestingly, in the case of donor sites, there was a higher frequency of SNPs neighboring events differentially spliced between Col-0 and L*er* accessions in the *prmt5-5* than in the WT background. This finding is consistent with PRMT5 assisting in the recognition of weak 5’ss, as it implies that changes in these sequences should have a greater impact in the absence of functional PRMT5. An example of a gene showing increased differential splicing between accessions in the *prmt5* compared to WT backgrounds, is shown in Fig. 3 C,D.

### PRMT5 effects are predicted by SNPs affecting donor splice strength

To better evaluate the impact of the genomic background on PRMT5 effects on splicing we focused our analysis on those splicing events for which the effect of the non-functional *PRMT5* allele on pre-mRNA splicing was different between accessions. We identified a total of 54 splicing events with a significant interaction effect between *PRMT5* allelic variants and the genomic background. At least one SNP in the donor splice site sequence was identified for 14 of the 54 events (Sup. Table 2). This value represented ∼25% of genes with significant allele*background interaction effects which is more than a two-fold enrichment compared to the number of genes with differentially spliced events detected between accessions regardless of the allele variant at the *PRMT5* locus (∼10%, see Fig. 3B). This enrichment strengthens the idea that PRMT5 plays a major role in modulating the impact that genetic variation at donor splice site sequences has on pre-mRNA splicing.

To further characterize the observed interaction, we evaluated the correlation between quantitative differences in PRMT5 effects on pre-mRNA splicing and the difference in donor splice site strength between accessions, which was quantified using a data-based energy model for donor sequences described in (Beckel et al. 2023). In that work, Beckel and collaborators introduced a Maximum Entropy Model to infer a meaningful energy scale that could assess splice site strengths from donor sequence compositions.

In Fig. 4A we show how differences in donor splice site strength were connected to the differences in IR between *prmt5-5* mutant and WT plants in the different accessions for the events showing allele*background interaction. As can be seen, in most cases where the *prmt5-5* mutation resulted in higher IR in L*er* relative to Col-0, this was associated with the 5’ss being weaker in L*er*. Similarly, in the cases where the effect of the *prmt5-5* mutation on IR was greater in Col-0, the weakest 5’ss variant was found in that accession.

**Figure 4.**
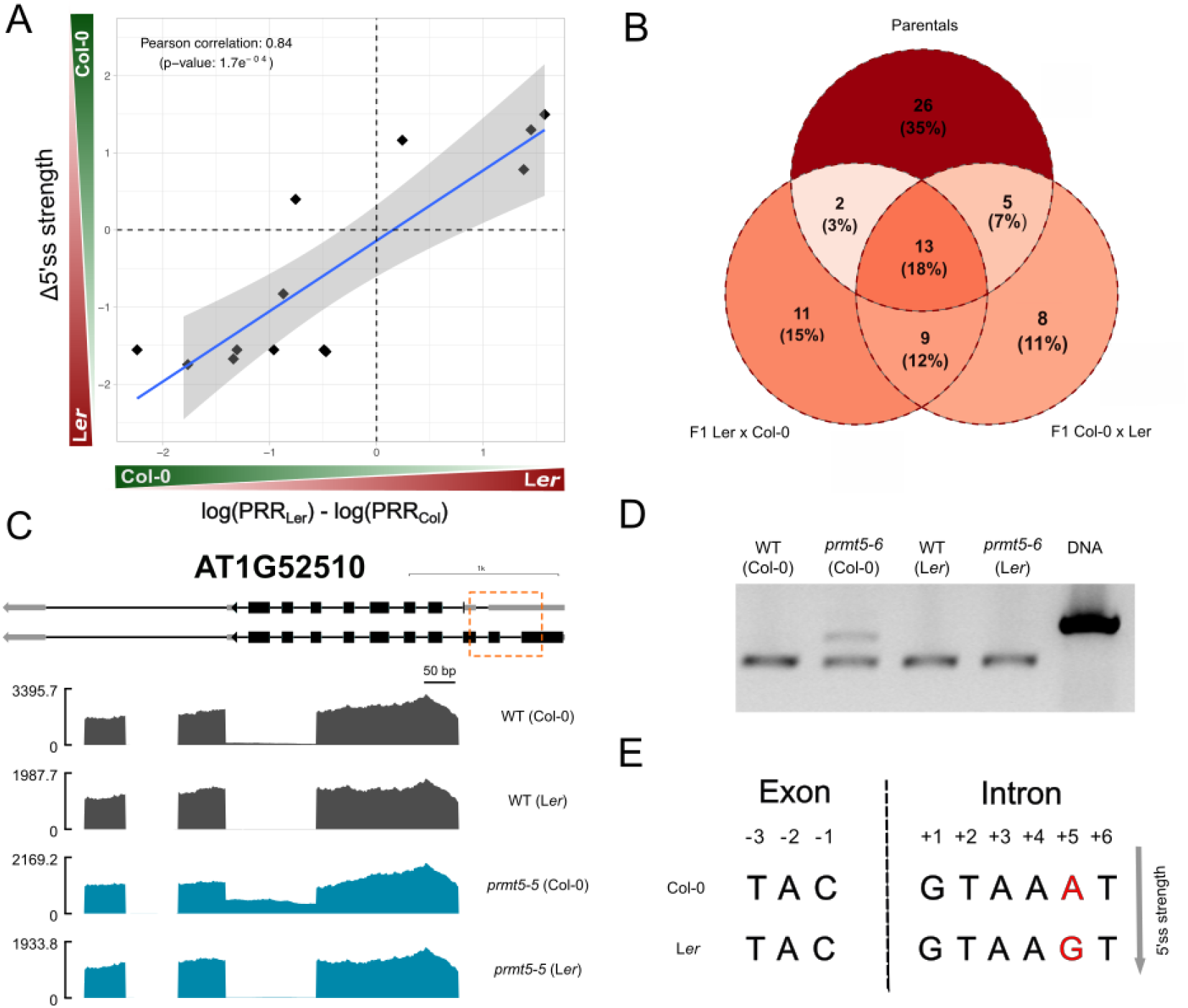
5’ss with sequence variation in genotype-accession interaction events. **(A)** Relationship between 5’ss strength differences produced by the presence of SNPs and the genotype x accession interaction effect on PIR, represented as the difference between L*er* and Col log-transformed PIR Relative Ratios (PRR=PIR_*prmt5-5* / PIR_WT). **(B)** Venn diagram of the significant interaction events in the F_1_ hybrids and in the subsamples of the parental plants. **(C)** Coverage plot of AT1G52510 in WT (gray) and *prmt5-5* (blue) Col-0 and L*er* accessions. A representative scheme describing the different isoforms annotated for the gene is shown above the coverage plot. Boxes represent exons, lines represent introns and the orange box surrounds the measured event. **(D)** Validation of the differential splicing event by agarose gel with the RT-PCR amplicons and the corresponding genomic DNA control. (**E**) Representative scheme of the 5’ss of the gene AT1G52510 in Col-0 and L*er*.

To better disentangle *cis* and *trans* contributions to the accession dependent effects of PRMT5 on splicing, we compared the effects of loss of *PRMT5* on pre-mRNA splicing in F_1_ hybrid progeny obtained by crossing Col-0 and L*er* accessions, with and without *prmt5* mutations. In the F_1_ hybrids, the global genetic environment is the same, differences in splicing between allelic variants can be attributed to changes in *cis*-sequences and not *trans*-regulatory factors.

Fig. 4B shows the results for the interaction effect analysis of the two types of reciprocal hybrids and the parental lines. We found that, out of a total of 46 events that were differentially spliced by PRMT5 between the parental plants, 43% were also found to be differentially spliced in an allele-dependent manner in the hybrids. Since in the F_1_ hybrids both alleles were in the same *trans* context, their differential response to the absence of functional PRMT5 was likely due to the differences in *cis* between the two accessions. We can therefore conclude that a significant fraction of the accession-dependent effects of PRMT5 on splicing were associated with genetic variation in *cis*. Furthermore, many of these effects were associated with SNPs affecting the strength of the donor splice sites associated with the splicing events affected.

### Impact of PRMT5 on phenotypic divergence between accessions

The effect of PRMT5 buffering the impact of genetic variation at donor splice sites on pre-mRNA splicing could contribute to PRMT5 acting as an evolutionary capacitor that, similarly to HSP90, suppresses phenotypic variation under normal conditions and releases this variation when functionally compromised. To evaluate whether PRMT5 could also be considered an evolutionary capacitor, we analyzed if it modulates the interplay between phenotypic and genotypic variation in *A. thaliana*. Interestingly, we found that PRMT5 significantly altered the range of phenotypic diversity in leaf morphology between Col-0 and L*er* accessions, as serrated leaves were observed in *prmt5* mutant plants in a Col-0 but not L*er* genomic background, while differences in leaf morphology were not that obvious between WT Col-0 and WT L*er* plants (Fig. 5A). A similar phenomenon was observed for flowering time, with differences in this trait being significantly larger between accessions in the *prmt5* mutant than in a WT background (Fig. 5B). A similar trend, although not statistically significant (P=0.06), was observed for the circadian period phenotype of the rhythm of leaf movement, with larger differences between accessions observed in the *prmt5* than in the WT backgrounds (Fig. 5C,D).

**Figure 5.**
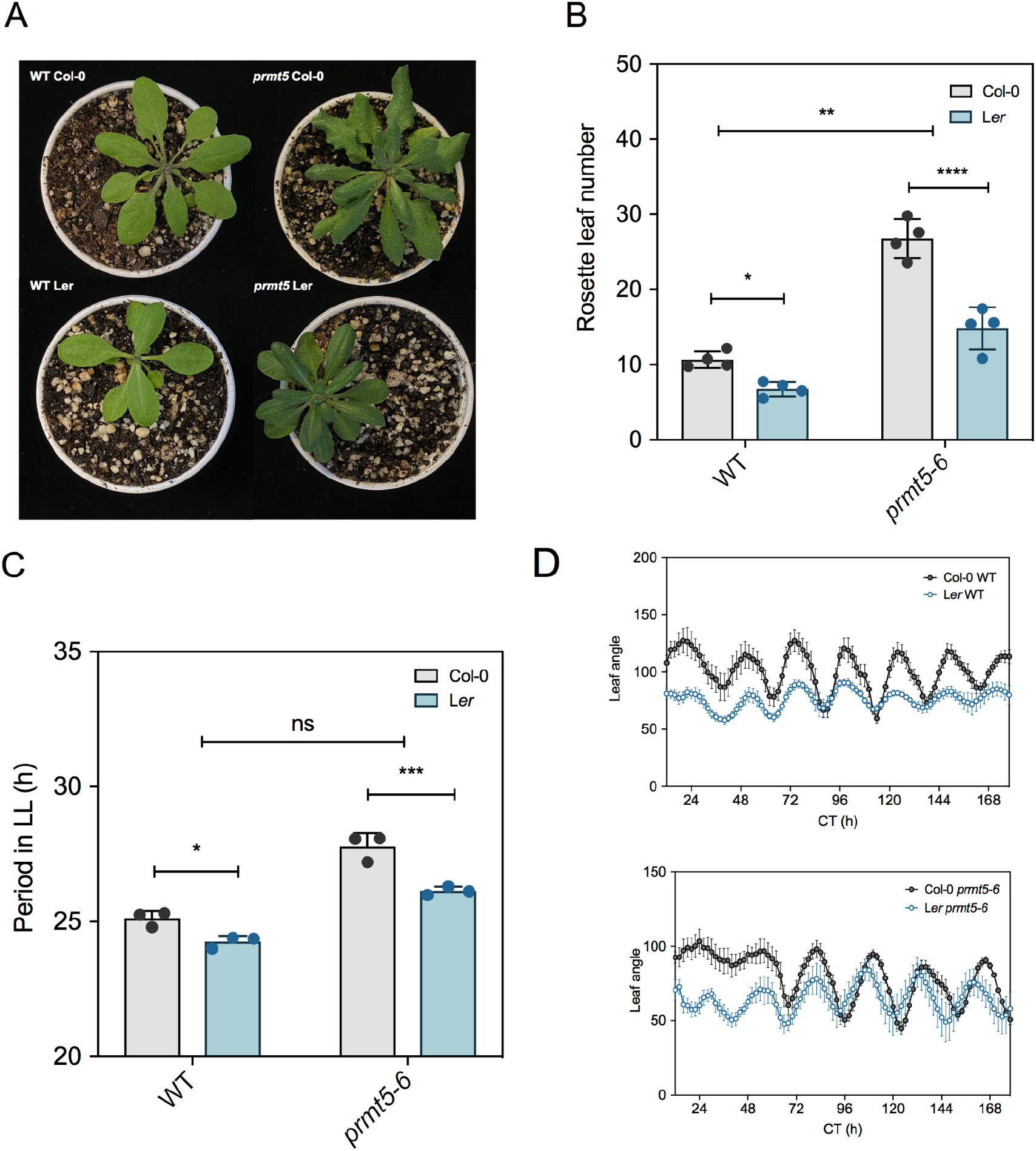
PRMT5 differentially affects physiological and developmental processes in the Col-0 and L*er* accessions. All the experiments were performed under a long-day photoperiod at 22°C. (**A**) Rosette phenotype at bolting of WT and *prmt5-6* mutant plants grown in soil in Col-0 and L*er* accessions. (**B**) Rosette leaf number at flowering time of WT and *prmt5-6* mutant in Col-0 and L*er* plants. (**C**) Periods of circadian rhythms in leaf movement calculated with BioDare2 for WT and *prmt5-6* mutant plants of the Col-0 and L*er* accessions. (**D**) Leaf angle of Col-0 and L*er* WT and *prmt5-6* in continuous light after entrainment under long-day conditions. Data represents mean + SEM (n≥ 3). Significant difference, as determined by two-way ANOVA followed by a Sidak’s test for multiple comparisons (* p< 0.05; **p < 0.01; *** p< 0.001; **** p< 0.0001; ns: not significant).

## DISCUSSION

### PRMT5 attenuates the effects of natural variation on splice site sequences

There is increasing evidence that the ability of organisms to adapt to environmental changes depends on pre-mRNA splicing, as it allows for the production of novel transcripts and/or protein isoforms from a single gene (Verta and Jacobs 2022). While genetic variation at splicing regulatory sequences may result in a novel isoform, it can also negatively impact pre-mRNA splicing and organismal function (Scotti and Swanson 2016). Therefore, the accumulation of cryptic genetic variants that remain phenotypically silent until circumstances change requires a buffering mechanism. The work described here suggests that PRMT5 acts as a master capacitor that buffers the effects of natural genetic variation on pre-mRNA splicing by enhancing the usage of weak donor splice sites present in the genetically diverse *A. thaliana* accessions Col-0 and L*er*.

The recent characterization of the divergence and regulatory mechanisms of AS among different accessions of *A. thaliana* revealed that sequence changes acting in *cis* were the major source of intraspecies variation in pre-mRNA splicing (X. Wang et al. 2019). Similar observations have been made in studies in mammals (Gao et al. 2015; Barbosa-Morais et al. 2012), suggesting that *cis*-regulatory divergence may be a common effect regulating intraspecies splicing divergence in plants and animals (X. Wang et al. 2019). The results presented here support these observations, revealing significant differences in pre-mRNA splicing patterns between L*er* and Col-0 accessions that were predominantly associated with genetic variation in nearby sequences. In addition, as expected, a strong purifying selection was found for mutations affecting the canonical +1 and +2 positions of the 5′ss, which are critical for the completion of the splicing reaction. Interestingly, the extent of divergent variation in pre-mRNA splicing between Col-0 and L*er* accessions was much larger in a *prmt5* mutant than in a WT background. Furthermore, we found that many of the divergent splicing events were enriched in those associated with sequence variants in donor-splice sites, with SNPs weakening splice sites leading to larger intron retention in *prmt5* mutants than in WT plants, while SNPs strengthening donor splice sites causing the opposite response. Finally, the analysis of F_1_ hybrid offspring supports the idea that sequence variation in *cis* associated with donor splice sites is significantly associated with the accession-dependent effects of PRMT5 on pre-mRNA splicing. Together, these observations suggest that the stabilizing effect of PRMT5 on pre-mRNA splicing is likely to play a relevant role in facilitating evolutionary adaptations by allowing genetic variation to persist without compromising gene function.

### Common and genotype-specific effects of PRMT5

PRMT5 plays a vital role in several interconnected processes, including transcription, mRNA processing and transport, and chromatin remodeling (Koh et al., 2015). This diversity of functions establishes PRMT5 as a crucial regulator of gene expression at multiple levels. Consistent with this, we identified a large common set of genes and splicing events associated with these processes that were similarly regulated by PRMT5 in both the Col-0 and L*er* accessions. Notably, we found that genes related to splicing and RNA metabolism were overexpressed in *prmt5* mutants compared to WT plants in both accessions, highlighting PRMT5’s key role in modulating these processes. Furthermore, as previously reported in both plants and animals, splicing events affected in *prmt5* mutants in either Col-0 or L*er* accessions were preferentially associated with weak donor splice sites (Sanchez et al. 2010; Bezzi et al. 2013; Hernando et al. 2015). Interestingly, in most cases for which a SNP in L*er* strengthened the 5′ss of a splicing event affected in the *prmt5* mutant in the Col-0 background, splicing of the event in the *prmt5* mutant in L*er* was much less affected, strongly supporting the idea that PRMT5 preferentially contributes to the recognition and usage of weak donor splice sites.

### Mechanisms associated with accession-dependent effects of PRMT5 on pre-mRNA splicing

PRMT5 may regulate splicing through multiple pathways, and the precise molecular mechanism of most of its effects on splicing is far from being understood. First, PRMT5 can enhance splicing efficiency by methylating and affecting the function of core and auxiliary splicing factors (Radzisheuskaya et al. 2019). Second, it may influence the expression of these factors through transcriptional regulation, partly via histone methylation (Koh, Bezzi, and Guccione 2015). This dual mechanism could compensate for reduced splicing efficiency due to lack of methylation of splicing factors, by increasing the expression of splicing components. However, it remains to be determined whether the expression changes observed in WT versus *prmt5* mutants in *A. thaliana* are linked to alterations in histone methylation or other transcription-regulating proteins. Furthermore, the precise reasons behind PRMT5’s preferential regulation of a subset of splicing events with weak 5’ss are still unknown. While most affected 5’ss are statistically weaker, not all weak sites are influenced by PRMT5. Analysis of divergent splicing between the Col-0 and L*er* backgrounds revealed that nearby sequence variations correlate with splicing changes. Although SNP density between accessions is identical in WT and *prmt5* mutant plants, a higher number of differential splicing events between accessions was observed in a *prmt5* mutant compared to a WT background. Notably, there was a greater occurrence of SNPs in the 5’ss regions of divergent events in the *prmt5* mutant background, suggesting that pre-mRNA splicing was more sensitive to genomic variations in donor splice sites in the absence of PRMT5. Furthermore, a significant correlation was found between variations in 5’ss strength and PRMT5’s impact on splicing efficiency evaluated as percentage intron retention, indicating that the donor splice site sequence is crucial for PRMT5’s role. These findings suggest that PRMT5 is essential for maintaining splicing efficiency amid genomic variations, potentially acting as a safeguard for system stability. Identifying nearby *cis*-regulatory sequences preferentially associated with weak donor splice sites whose usage depends on PRMT5 activity could eventually contribute to a better understanding of the particular mechanism(s) through which PRMT5 regulates the splicing of specific events. Most likely, PRMT5 methylates and regulates the activity of one or more splicing factors that modulate the splicing efficiency of nearby associated introns, but the identify of such factors remains unknown.

A large fraction of the events detected in the individual accessions were also observed in their F_1_ hybrid progeny. However, to fully comprehend these findings, it is important to note the methodological limitations. The sequencing of hybrid mRNA was conducted in bulk, and each read was attributed to one of the two accessions based on the genome that achieved higher mapping quality. Reads that gained identical mapping quality for both accessions’ genomes were omitted from the analysis. Consequently, genome areas with low SNP/Indel density between Col-0 and L*er* endured a significant drop in coverage. This inhibits them from being statistically evaluated. Upon analysis of the events that had a significant interaction effect on the parental plants but not on the hybrids, it was found that these regions of the genome have a decreased density of sequence variations. Based on these considerations, it is reasonable to assume that the observed percentage of coincidence between the events of the parents and the hybrids might underestimate the true level of coincidence if the density of variations were uniform across different genes. This further supports the notion that PRMT5 effects are strongly impacted by the *cis*-sequences located near the splicing events. A more in-depth understanding of the *cis-trans* interactions associated with PRMT5 effects on splicing is likely to be obtained by future analysis of the transcriptomes of parental plants and hybrids with long-read sequencing technologies, which allow for the unambiguous assignment of each read to a specific accession.

Finally, our findings, together with the results from other groups, provide additional considerations that need to be taken into account for a full understanding of how PRMT5 affects splicing patterns. Recent studies indicate that PRMTs selectively impact post-transcriptional splicing (Maron et al. 2022; Yan et al. 2024), which, while comprising only about 20% of splicing events in human cells and 28% in *A. thaliana*, plays a crucial physiological role (Jia et al. 2020; Girard et al. 2012). Transcripts with unremoved introns remain in the nucleus until a signal triggers their removal, allowing for cytoplasmic export. This mechanism is advantageous during stress, as it speeds up responses and protects these transcripts from degradation via nonsense-mediated decay (NMD). Post-transcriptional splicing is often associated with nuclear speckles, which serve as reservoirs for splicing factors and are active sites for splicing. Various post-translational modifications, including arginine methylation, regulate the structure of these speckles. Splicing events processed post-transcriptionally tend to occur at weak splice sites, a characteristic also seen in PRMT5-dependent events. A deeper understanding of the mechanisms affecting post-transcriptional splicing will therefore contribute to the improvement of our understanding of PRMT5 effects on pre-mRNA splicing and their dependence on 5’ ss strength.

### Evolutionary Implications

Buffering and compensatory mechanisms are essential for managing the effects of natural genetic variation on gene expression and phenotype. By ensuring stability and consistency in phenotypic traits across genetic and environmental perturbations, a phenomenon known as canalization, these processes contribute significantly to the resilience and adaptability of living organisms (Rutherford 2000). HSP90 is an example of a regulatory protein that plays a significant role in canalization, acting as a capacitor that buffers the effects of genetic variation on phenotypic expression (Rutherford and Lindquist 1998; Queitsch, Sangster, and Lindquist 2002). Key properties of HSP90 that led to its proposal as an evolutionary capacitor are that HSP90 suppresses phenotypic variation under normal conditions and releases this variation when functionally compromised; its biological functions are associated with and are affected by, responses to environmental stress and, finally, it exerts pleiotropic effects on key developmental processes (Bergman and Siegal 2003). Similarly to HSP90, PRMT5 has pleiotropic phenotypic effects, and its functions have been associated with plant responses to environmental stress. Furthermore, the loss of PRMT5 significantly enhanced differences in splicing patterns between accessions and this was associated with larger phenotypic variation in developmental processes. Therefore, PRMT5 appears to fulfill the criteria for being considered a capacitor that buffers the effects of natural genetic variation on pre-mRNA splicing and developmental phenotypes.

The buffering role of PRMT5 on pre-mRNA splicing has implications for evolutionary biology. By masking the effects of many splice site polymorphisms, PRMT5 facilitates the exploration of new splicing patterns that could confer adaptive advantages under changing environmental conditions. This capacity for phenotypic plasticity is essential for survival and reproduction, particularly in fluctuating ecosystems. Furthermore, the study raises important questions about the evolutionary conservation of PRMT5 function across species. Given that AS is a key driver of diversity in both plants and animals, understanding how similar mechanisms operate in different organisms could provide insights into evolutionary processes at large. Further research is needed to explore the conservation of PRMT5’s buffering role on pre-mRNA splicing and its potential clinical relevance in humans, where PRMT5 inhibitors are increasingly being used as a treatment for different types of cancers.

## CONCLUSION

AS plays a significant role in evolutionary processes. Variations at core and regulatory splice sites can lead to alternative splicing, generating diverse transcript and protein isoforms that introduce phenotypic variability, which canalization mechanisms should buffer. Our findings highlight how the interaction between sequence variants at donor splice sites and PRMT5 modulates the effects of weak donor splice sites on pre-mRNA splicing, demonstrating that PRMT5 can help populations maintain splicing stability while harboring latent diversity. When environmental conditions shift, these previously masked variants can contribute to phenotypic diversity, enabling evolutionary adaptations.

## ACKNOWLEDGEMENTS

This work was supported by grants from Agencia Nacional de Promociones Científicas y Tecnológicas (MJY, ACh) and the Max Planck Society (DW). We thank the members of our labs for their critical reading of the manuscript.

