## Supplemental Material for "Arabidopsis PRMT5 Buffers Pre-mRNA Splicing and Development Against Genetic Variation in Donor Splice Sites"

A.

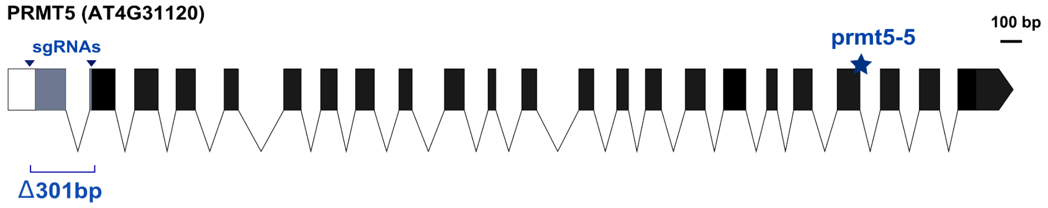


B.

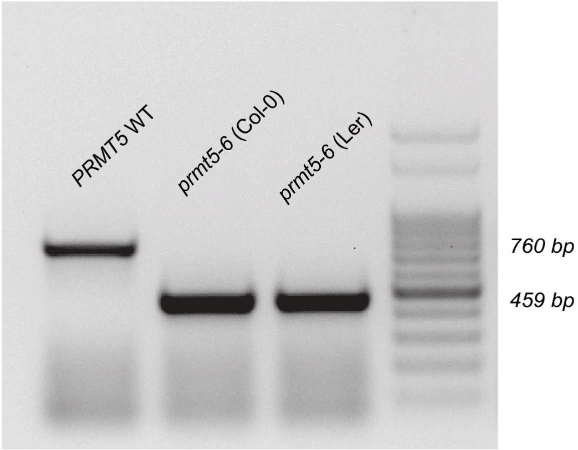


**Supplementary Figure S1.** (**A**) Schematic representation of the *prmt5* mutations. The exons are indicated by black boxes, and the introns are represented by black lines. The position of the *prmt5-5* point mutation is indicated by a star, and the single guide RNAs (sgRNAs) utilized for CRISPR-mediated edition are represented by triangles. The *prmt5-5* mutation consists of a single base exchange that introduces a premature stop codon. The CRISPR mutation produces a deletion of 301 bp, which is indicated by the gray box in the scheme. (**B**) Agarose gel electrophoresis of the PRMT5 WT gene and CRISPR *prmt5-6* mutant in Col-0 and Ler accessions. The numbers indicate the weight of the amplicon in base pairs (100 bp DNA ladder).

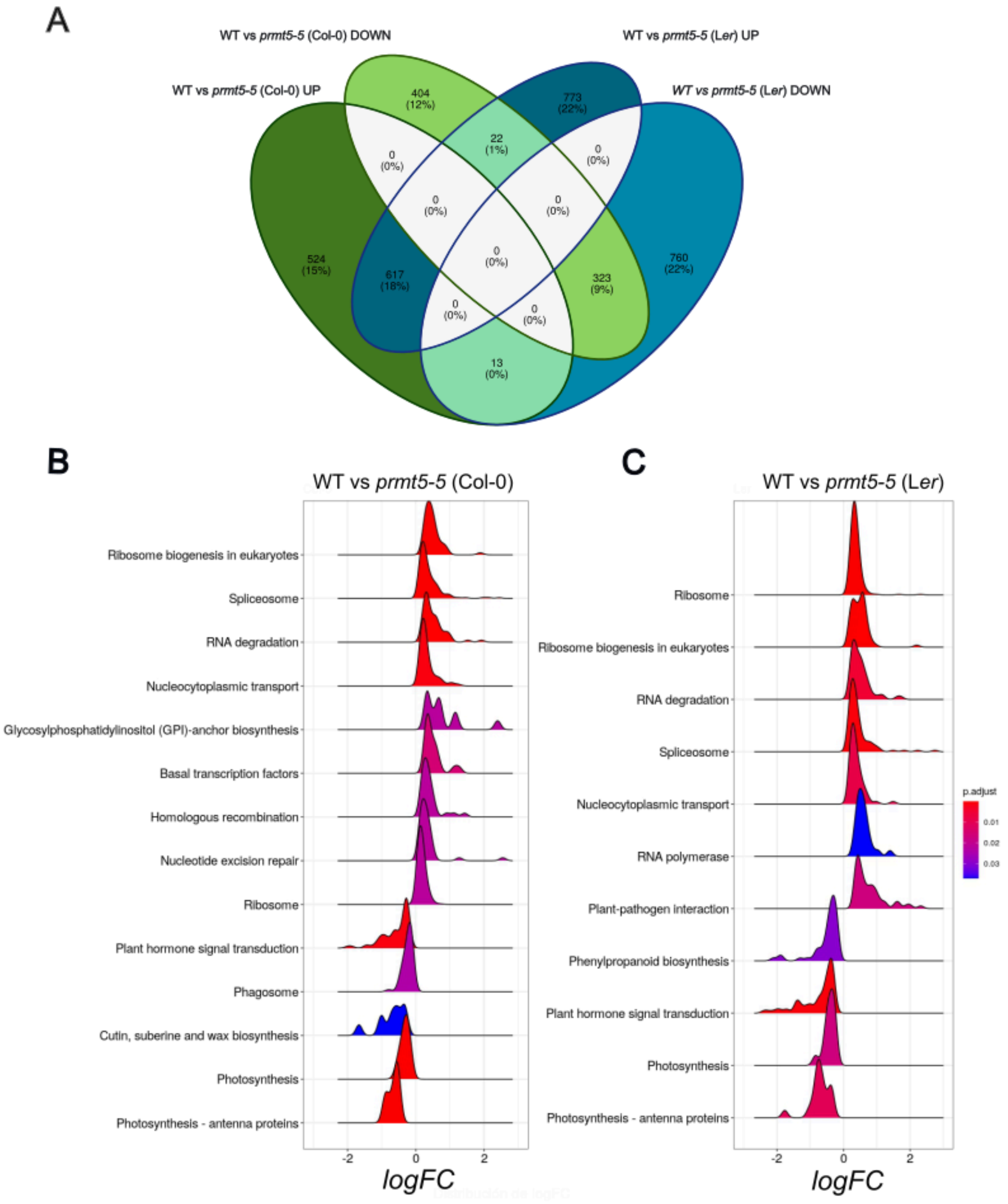


**Supplementary Figure S2.** Differentially expressed genes (DEG). Comparison between WT and mutant plants for PRMT5 in both accessions, Col-0 and L*er*. (**A**) Venn diagram of the DEGs found in the analysis. It shows the sets of genes whose expression was significantly higher in the prmt5-5 mutant (Col-0 Up and L*er* Up) and those whose expression was found to be lower (Col-0 Down and L*er* Down). LogFC distribution of genes whose pathways were found to be significant in the KEGG enrichment analysis for WT vs *prmt5-5* in Col-0 accession (**B**) and L*er* accession (**C**).

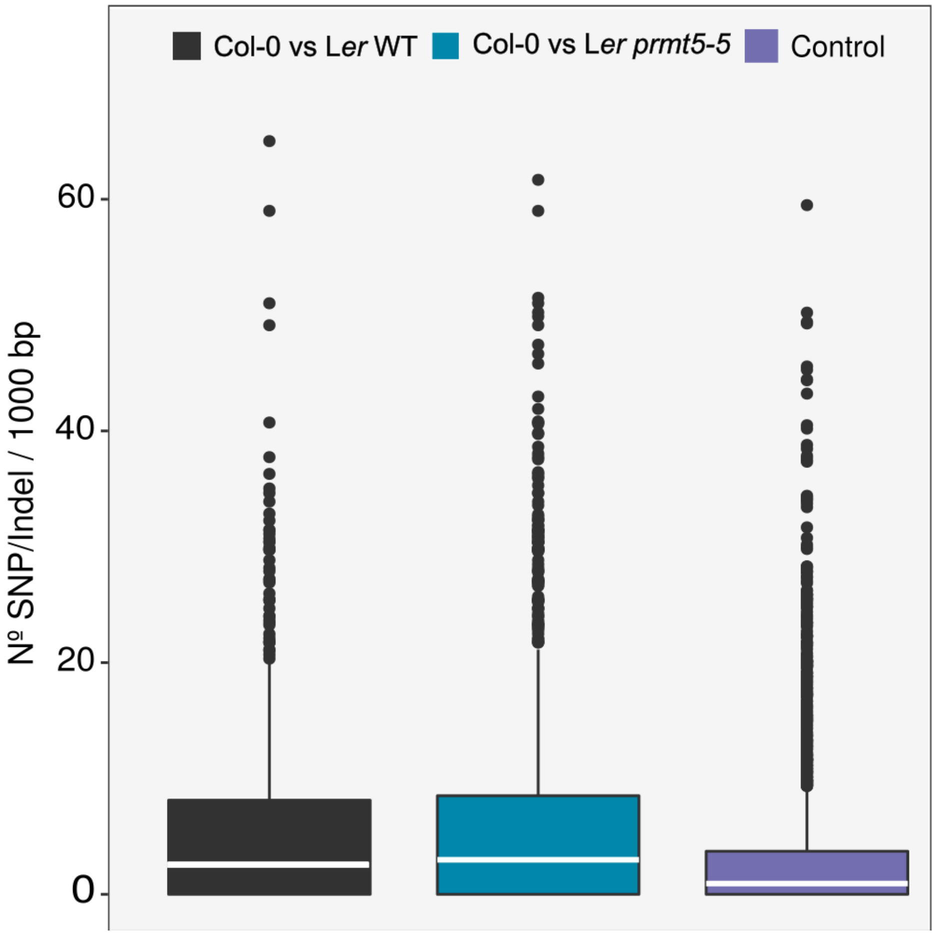


**Supplementary Figure S3.** Density of sequence variation between Col-0 and L*er* around IR events in WT and *prmt5-5* and for IR events annotated in the reference genome but did not give differential signal (Control).

**Supplementary Table S1:** All the primers used in this work

| Primer name | Sequence (5'-3') | Use |
|---|---|---|
| DT1_BsF_PRMT5 | ATATATGGTCTCGATTGtctctctctattgagctcacGTT | for CRISPR |
| DT1_F0_PRMT5 | TGtctctctctattgagctcacGTTTTAGAGCTAGAAATAGC | for CRISPR |
| DT2_BsR_PRMT5 | AACccaagctcggcctatacgaaCAATCTCTTAGTCGACTCTAC | for CRISPR |
| DT2_R0_PRMT5 | ATTATTGGTCTCGAAACccaagctcggcctatacgaaCAA | for CRISPR |
| prmt5_crispr_check_Fw | TGGGATGGCATAAATGGGCTAGG | for check mutation |
| prmt5_crispr_check_Rv | AAGGAGAGATGAGTAGCCCAAGC | for check mutation |
| CAS9_Fw | CATATCGTTCCACAGTCATTCC | for check Cas9 in pHEE401E |
| CAS9_Rv | GGTAGCCTTGCCAATCTCC | for check Cas9 in pHEE401E |
| AT3G61420_IR_Fw | AGGACGCTAATCAGGAGAGG | AS event validation |
| AT3G61420_IR_Rv | TTGGCAGTTGAGATAGTTCGG | AS event validation |
| AT4G28470_IR_Fw | GGTTGCTATGTCTGAGGAATTGG | AS event validation |
| AT4G28470_IR_Rv | GCACTGGCACCGAGATGG | AS event validation |
| AT1G52510_IR_Fw | CTTCCTCTGCTCCACTCATTCTCC | AS event validation |
| AT1G52510_IR_Rv | CAATGGTTCCTCGACGACTCTCC | AS event validation |
| AT5G02810_IR3_Fw | AGAGTCGGACAGAGAAGCATAG | AS event validation |
| AT5G02810_IR3_Rv | CCGCTCTCACTTCCACTACC | AS event validation |

**Supplementary Table S2**: Interaction splicing events between allele and background with at least one SNP in the donor splice site sequence. **Locus** indicates the name of the gene; **PRR_Col** and **PRR_L*er*** indicate the PIR relative ratio between *prmt5* and WT estimated for Col and L*er* accessions respectively; **p-value** accounts for the statistical significance of the difference between log-transformed **PRR_L*er*** and **PRR_Col** values; **FDR**: False Discovery Rate; **seq_5ss_Col** and **seq_5ss_L*er*** indicate the sequence of the donor 5' splice site in Col and L*er* accessions, respectively, and **e_5ss_Col** and **e_5ss_L*er*** indicate the energy of these 5's in Col and L*er* accessions, respectively.

| locus | PRR_ Col | PRR_ Ler | p-value | FDR | seq_5ssCol | seq_5ssLer | e_5ss_ Col | e_5ss_ Ler |
|---|---|---|---|---|---|---|---|---|
| AT1G44446 | 2,68 | 1,67 | 5,82E-03 | 3,47E-02 | AAGGTTTCT | AAGGTTTGT | 7,312 | 5,733 |
| AT1G52510 | 9,33 | 1,59 | 3,06E-03 | 2,07E-02 | TACGTAAAT | TACGTAAGT | 10,238 | 8,493 |
| AT1G65540 | 4,68 | 1,27 | 1,92E-02 | 8,83E-02 | GGAGTTAGT | GGAGTAAGT | 9,432 | 7,877 |
| AT1G69620 | 18,68 | 1,98 | 6,14E-28 | 4,99E-26 | GGTGTTAGT | GGTGTAAGT | 9,359 | 7,804 |
| AT3G14660 | 5,92 | 2,77 | 9,64E-31 | 1,18E-28 | AAAGTTTAT | AAAGTTTAA | 9,762 | 10,159 |
| AT3G55630 | 2,74 | 1,14 | 1,91E-02 | 8,83E-02 | AAGGTGTTT | AAGGTGTGT | 7,358 | 6,53 |
| AT4G12460 | 1,24 | 5,99 | 1,98E-03 | 1,46E-02 | GAGGTATAT | GAGGTAGAT | 6,183 | 7,68 |
| AT4G16765 | 0,96 | 3,79 | 8,70E-05 | 1,18E-03 | CAGGTGTGT | CAGGTTTCT | 6,634 | 7,417 |
| AT4G17460 | 2,01 | 0,77 | 1,98E-02 | 8,97E-02 | CCCGTTAGT | CCCGTAAGT | 9,453 | 7,898 |
| AT4G25640 | 2,19 | 1,34 | 1,18E-02 | 6,40E-02 | ATGGTTAGA | ATGGTAAGA | 8,36 | 6,805 |
| AT5G02810 | 1,09 | 1,39 | 4,16E-03 | 2,57E-02 | TCAGTGAGT | TCAGTGAGA | 8,513 | 9,675 |
| AT5G07440 | 3,66 | 1,13 | 1,40E-03 | 1,14E-02 | CAAGTAAAA | CAGGTAAAA | 8,189 | 5,57 |
| AT5G11840 | 11,73 | 3,07 | 1,62E-03 | 1,27E-02 | AGTGTACTT | AGTGTAATT | 11,07 | 9,397 |
| AT5G59480 | 0,56 | 2,37 | 9,31E-03 | 5,41E-02 | TCGGTATGT | TCTGTATGT | 7,108 | 8,406 |
